# Simultaneous profiling of DNA methylation and chromatin architecture in mixed populations and in single cells

**DOI:** 10.1101/470963

**Authors:** Guoqiang Li, Yaping Liu, Yanxiao Zhang, Rongxin Fang, Manolis Kellis, Bing Ren

**Affiliations:** Ludwig Institute for Cancer Research, La Jolla, CA 92093; Department of Cellular and Molecular Medicine, Center for Epigenomics, Institute of Genomic Medicine, Moores Cancer Center, University of California, San Diego, School of Medicine, La Jolla, CA 92093; Broad Institute of MIT and Harvard, Cambridge, MA 02142; Massachusetts Institute of Technology, Computer Science and Artificial Intelligence Laboratory, Cambridge, MA 02139; Bioinformatics and Systems Biology Graduate Program, University of California, San Diego, La Jolla, CA 92093

## Abstract

Dynamic DNA methylation and three-dimensional chromatin architecture compose a major portion of a cell’s epigenome and play an essential role in tissue specific gene expression programs. Currently, DNA methylation and chromatin organization are generally profiled in separate assays. Here, we report Methyl-HiC, a method combining *in situ* Hi-C and whole genome bisulfite sequencing (WGBS) to simultaneously capture chromosome conformation and DNA methylome in a single assay. Methyl-HiC analysis of mouse embryonic stem cells reveals coordinated DNA methylation between distant yet spatially proximal genomic regions. Extension of Methyl-HiC to single cells further enables delineation of the heterogeneity of both chromosomal conformation and DNA methylation in a mixed cell population, and uncovers increased dynamics of chromatin contacts and decreased stochasticity in DNA methylation in genomic regions that replicate early during cell cycle.

---

DNA methylation plays a critical role in gene regulation^1^. DNA methylation is dynamically regulated by a variety of enzymes, including the *do novo* methyltransferases DNMT3a and DNMT3b and the ten-eleven translocation (TET) family of dioxygenases TET1/2/3^2^. Additionally, DNA methylation patterns are maintained by DNMT1^3^. The methylation status of adjacent CpG dinucleotides on the same DNA fragment is often coordinated, a phenomenon that has been used to define methylation haplotype in liquid biopsy tests^4,5^. However, due to the short fragment length in Whole Genome Bisulfite Sequencing (WGBS)^6^ and absence of long-read methylome sequencing technologies^7,8^, it is not clear how long such coordinated DNA methylation extends in mammalian cells. As the DNA is spatially organized into three-dimensional structures, genomic regions that reside up to hundreds of kilobases away can be brought into spatial proximity through chromatin folding^9^. Thus, it is conceivable that DNA methylation between distal sequences could also be coordinated due to their spatial proximity. The chromosome conformation capture (3C) technology is based on proximal ligation of spatially close genomic DNA segments^10,11^. Since DNA methylation is a covalent modification on DNA, methylation status of cytosines far apart on the linear sequence could in principle be captured simultaneously on ligated DNA during chromosome conformation capture procedures. We exploit this principle to develop Methyl-HiC, combining *in situ* Hi-C^12^ and WGBS to simultaneously profile chromatin organization and DNA methylation genome-wide (Fig. 1a). Briefly, long-range chromatin interactions are captured by crosslinking with formaldehyde, digested with methylation insensitive restriction enzyme DpnII, labeled with biotinylated nucleotides, and ligated *in situ*. The ligation products are then enriched with streptavidin coated magnetic beads after sonication of genomic DNA. The captured DNA is then subject to bisulfite conversion, library construction and paired-end sequencing (Fig. 1a). We developed a computational pipeline, Bhmem, to map the sequencing reads to the reference genome, reveal the methylation status on linked DNA fragments (Fig. 1b and Supplementary Fig. 1a) (see Method for description of Bhmem) and compute the pairwise contact frequency genome wide.

**Fig.1.**
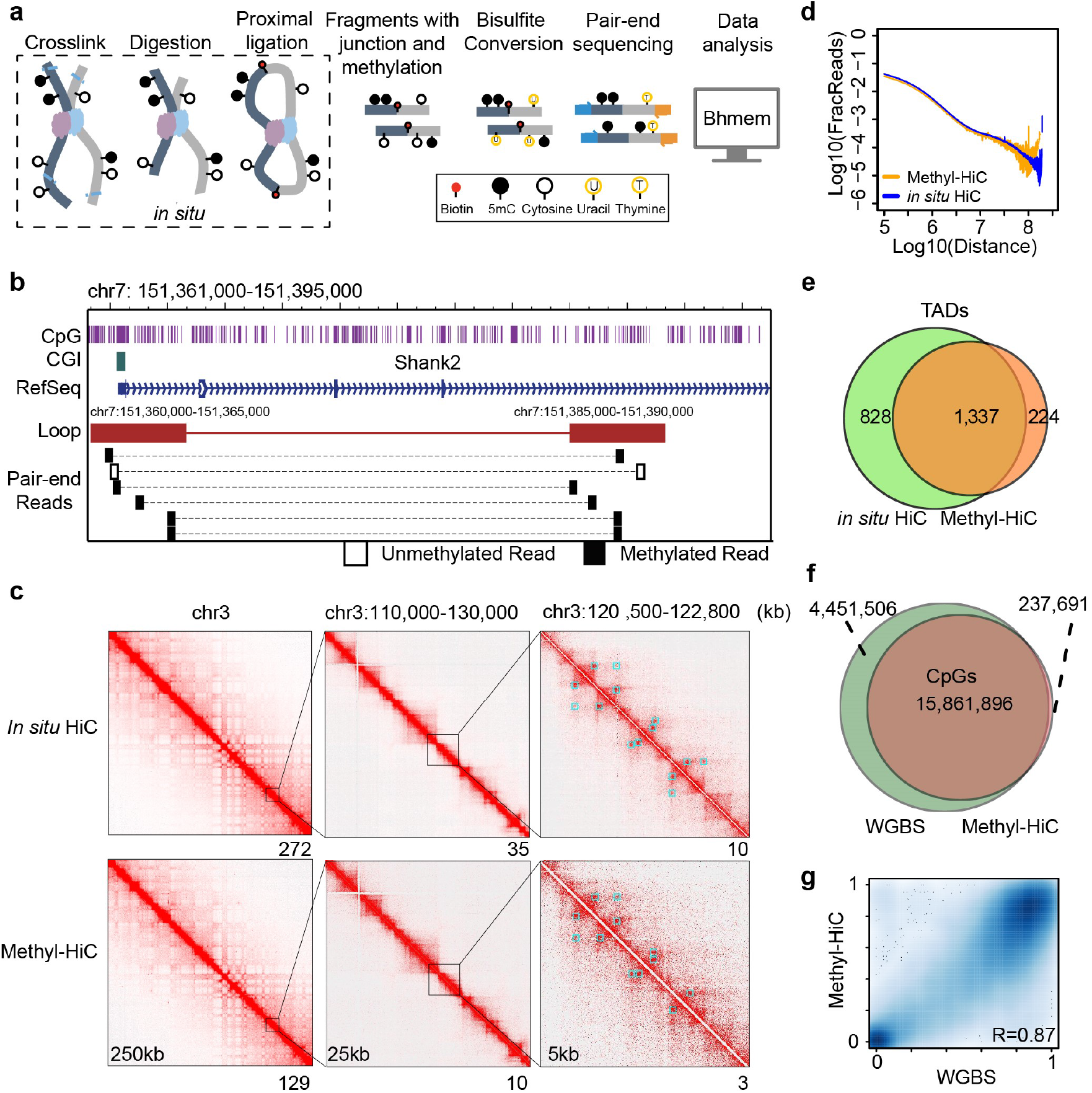
Methyl-HiC simultaneously profiles long-range chromatin interactions and DNA methylome in mouse embryonic stem cells. a) The workflow of Methyl-HiC. Biotin-enriched DNA fragments from *in situ* Hi-C are bisulfite converted followed by paired-end sequencing. The short sequencing reads are mapped to the genome and processed by the in-house computational pipeline Bhmem. b) An illustration shows the methylation status on read pairs supporting a chromatin loop. Methylation status are determined by CpG sites on the reads. c) Comparison of contact matrix between *in situ* Hi-C and Methyl-HiC at different resolutions. Blue squares are loops identified from corresponding dataset at 5kb resolution. Numbers below each map show the maximum values for the map color range. d) Comparison of contact frequency distance decay curve obtained from *in situ* Hi-C and Methyl-HiC data. e) Comparison of TADs identified from *in situ* Hi-C and Methyl-HiC. f) Comparison of CpG sites covered by at least 10 reads in WGBS and Methyl-HiC. g) DNA methylation levels for common CpGs are highly concordant between WGBS and Methyl-HiC.

To demonstrate the performance of Methyl-HiC, we first applied it to the mouse embryonic stem cell line F123, a hybrid between the castaneus and S129/SvJae mouse strains. We compared the results with that from *in situ* Hi-C analysis from the same cell line. With similar sequencing depth, Methyl-HiC results are highly correlated with *in situ* Hi-C at different resolutions (Fig. 1c, stratum-adjusted correlation coefficient (SCC)^13^ in 250kb resolution: 0.92±0.02; SCC in 25kb resolution: 0.88±0.008). The contact probabilities of two datasets are comparable to each other (Fig. 1d). The chromatin loops, detected using HiCCUPS^12^ at different resolutions, largely overlap between the two datasets (Fig. 1c, blue squares, and Supplementary Table 1). Additionally, chromatin loops identified from Methyl-HiC results showed similar enrichment of enhancers and promoters as marked by histone markers (H3K4me1, H3K27Ac, and H3K4me3), CTCF, and Polycomb-Repressed chromatin (H3k27me3), as *in situ* Hi-C, indicating that Methyl-HiC can effectively detect chromatin loops (Supplementary Fig. 1b). Furthermore, the Topologically Associating Domains (TADs) identified from the two datasets are also similar (Fig. 1e and Supplementary Fig. 1c). These results show that Methyl-HiC can capture chromosomal conformation as effectively as *in situ* Hi-C.

In addition to efficiently capturing chromosomal architecture, Methyl-HiC also profiles DNA methylation genome wide. We compared Methyl-HiC data with WGBS data from the same cell line and found that Methyl-HiC can capture about 80% CpGs from WGBS data (Fig. 1f). The methylation level of these CpGs showed great concordance (R=0.87, p<2.2-e16) with WGBS results (Fig. 1g and Supplementary Fig. 1d). We also notice that Methyl-HiC reads tend to be enriched at regions with DNA hypomethylation. Despite this bias, Methyl-HiC accurately measures the DNA methylation state for the over 15 million CpGs in the mouse genome (Fig.1g). These results, taken together, demonstrate that Methyl-HiC can simultaneously map DNA methylome and chromatin architectures in a biological sample.

Previous studies have reported that adjacent CpGs usually share concordant methylation status, and the stretches of DNA that contain such CpGs are termed methylation haplotype blocks^4,5^. Because the genome is not only linearly separated but also spatially organized, we hypothesize that spatially proximal DNA may also have coordinated methylation status. To test this hypothesis, we analyzed the chromatin loops detected from Methyl-HiC and *in situ* Hi-C in the same cell type. Indeed, the Pearson correlation coefficients of the methylation status from Methyl-HiC read pairs mapped to anchors of these chromatin loops (Supplementary Fig. 2a) are significantly higher than that from shuffled read pairs mapped to the same anchor regions (Fig. 2a and Supplementary Fig. 2b) (*p*<2.2e-16, Fisher z-transformation), indicating that methylation status of spatially proximal CpGs is coordinated. We further classified the loops according to the chromatin compartments they belong to. We observed that loop anchors in compartment A had higher methylation concordance compared to these in compartment B (Fig. 2b). This is in line with recent reports that the methylation correlation signal is a better predictor of A/B compartments than the average methylation signal^14^.

**Fig.2.**
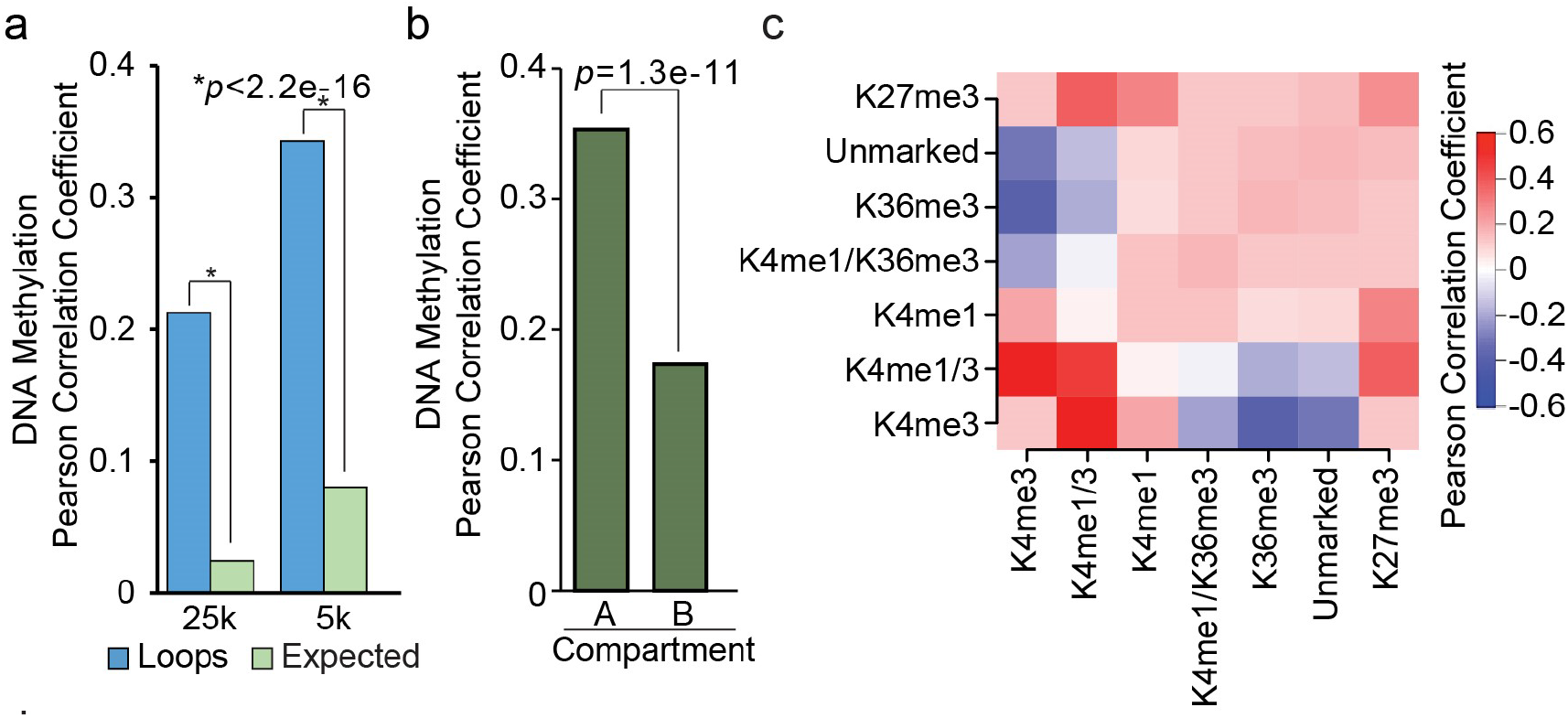
DNA methylation status is generally concordant between spatially proximal regions. a) Pearson correlation coefficient of DNA methylation concordance between anchors of chromatin loops identified using HiCCUPS at 5kb and 25kb resolutions. Only reads containing 2 or more CpGs on each end were included. Paired reads located within the same loop anchors were re-shuffled and used to calculate the expected values. b) Loops in compartment A show significantly higher concordance compared to those in compartment B. c) Concordance of DNA methylation levels between regions bearing various chromatin states called by chromHMM. Pearson correlation coefficients of DNA methylation were calculated from reads that fall within any two chromatin states.

Active and poised enhancers frequently exhibit hypomethylation in cells, and it is not clear whether the variable DNA methylation at enhancers is correlated with target gene promoters or other enhancers in the same transcription hub. Methyl-HiC data provides an opportunity to address this question. We classified the F123 cell genome into 7 different chromatin states using ChromHMM analysis of four histone modifications (H3K4me3, H3K4me1, H3K36me3, and H3K27me3)^15^. Pearson correlation coefficients of DNA methylation on the same Methyl-HiC read pairs showed different trends for pairs of genomic regions from different chromatin states. Methylation of DNA at actively transcribed genes, which are generally marked by H3K36me3, shows negative correlation with methylation status of active promoters and enhancers (Fig. 2c), consistent with previous observation that DNA methylation on TSS and gene body are inversely correlated^16^. By contrast, DNA methylation from active enhancer-like regions, marked by H34K4me1/3, shows positive correlation with that of linked promoters marked by H3K4me3, supporting the coordinated DNA methylation processes between active enhancers and their target gene promoters. Surprisingly, DNA methylation levels at regions with active enhancer-like chromatin state also display a positive correlation with that at linked regions marked by Polycomb-repressed chromatin state (Fig. 2c). This finding raises interesting questions regarding the relationships among polycomb-repressed state and enhancers, which will require additional experiments to resolve in the future.

Methods to map DNA methylome in single cells have been developed^17–19^, and used to deconvolute sub-populations from heterogeneous tissues or cell populations. Similarly, approaches to map chromatin organization in single cells have also been devised to reveal cell to cell variations in chromosome conformation in a mixed cell population^20–22^, and to study chromatin architecture in cell cycle^23^ and rare cell types, such as oocytes and zygotes^24^. One drawback of these methods is that DNA materials are destroyed during the process, preventing analysis of different epigenomic features from the same cells. The development of Methyl-HiC would overcome this limitation and provide the opportunity to explore the heterogeneity of DNA methylation and chromatin architecture simultaneously in a mixed cell population. We therefore modified methyl-HiC protocols to single cells. Briefly, after *in situ* proximal ligation, we sort individual nuclei into a 96-well plate, and then perform bisulfite conversion in each well. DNA adaptors are next ligated to the resulting bisulfite-converted single strand DNA (ssDNA), and the resulting DNA is PCR amplified for paired-end sequencing^17^ (Fig. 3a). We generated single cell Methyl-HiC data for 108 mES cells cultured in serum plus LIF condition (primed state) and 48 mESCs cultured in 2i conditions (naïve state)^25^. After removing low quality reads (mapQ<30), PCR duplicates, and inter-chromosomal read pairs, and cells with less than 250,000 reads and 10,000 *cis* ligations, we obtained data for 103 primed state mESCs and 47 naïve mESCs, each with about 1 million uniquely mapped and high quality intra-chromosomal reads for further analysis (Supplementary Table 2). Reads with each end originating from distinct restriction fragments are selected for further analysis, resulting in around 100,000 contacts on average in each cell (Supplementary Fig. 3a). The contact probabilities are comparable between the bulk dataset and the aggregate of single cells (Supplementary Fig. 3b). After normalization for sequencing coverage^26^, the intra-chromosomal interaction matrix from the bulk dataset and ensemble dataset of 103 primed mESCs are very comparable (Fig. 3b and Supplementary Fig. 3d, 3e), with the Pearson correlation coefficient computed from observed over expected contact matrix (250kb bin resolution) at 0.98 (*p*<2.2e-16). We also generated single cell methylomes from these cells with an average coverage of 288,000 CpGs per cell, a level that is comparable to previous single cell Methylome datasets^17,18^ (Supplementary Fig. 3c). The average methylation levels of primed and naïve mESCs are around 60% and 20% (Fig. 3c), respectively, consistent with previous single cell DNA methylome data^18^. Moreover, consistent with the observation from bulk populations, DNA methylation levels at the loop anchors are coordinated in individual cells (Fig. 3d). Taken together, these results show that our single cell Methyl-HiC can capture DNA methylation and chromatin architecture simultaneously in single cells.

**Fig.3.**
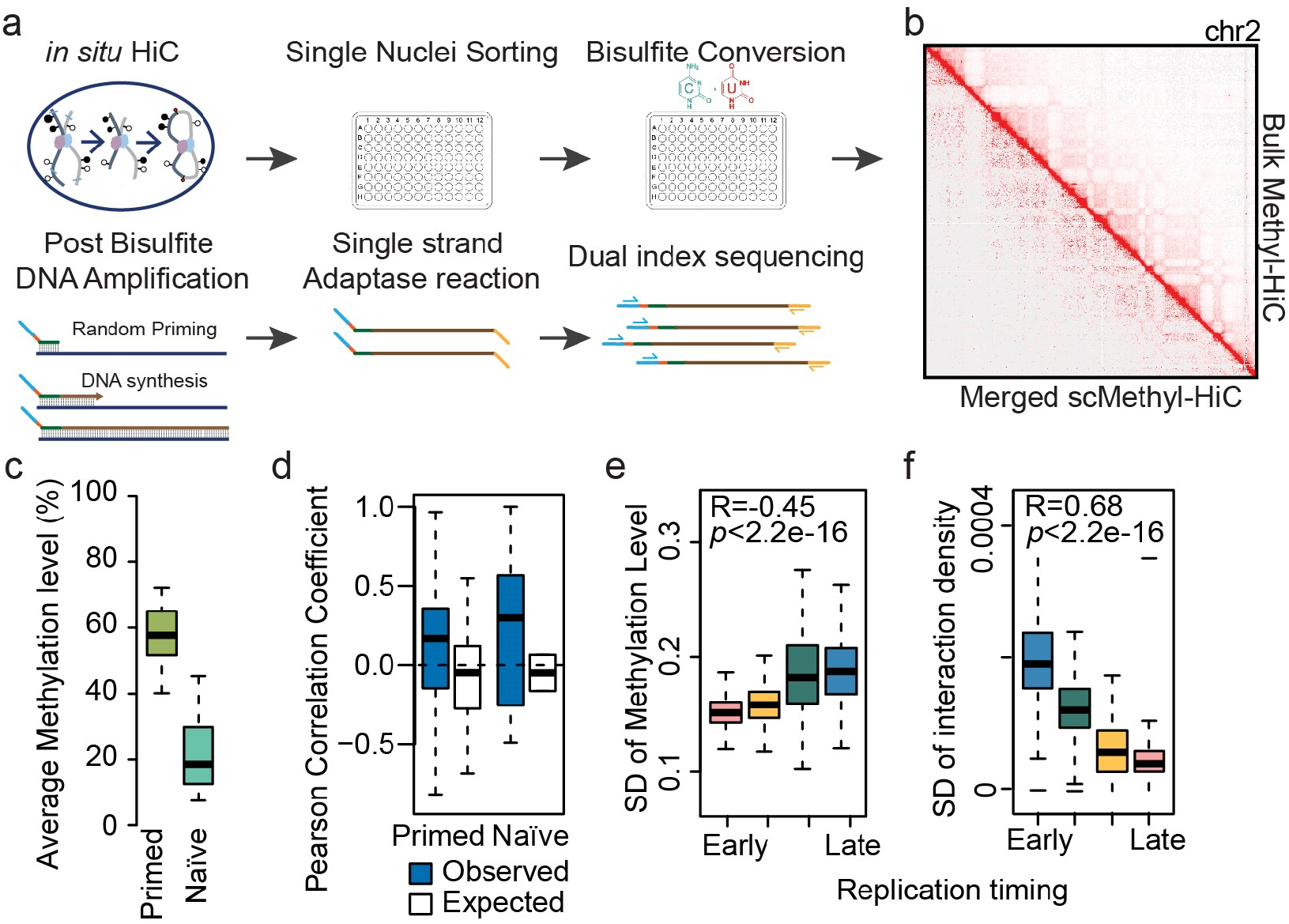
Simultaneous analysis of DNA methylome and chromatin architecture in individual cells by Single Cell Methyl-HiC. a) Workflow of Single cell Methyl-HiC. b) Comparison of aggregate contact matrix from 103 primed mESCs single cell Methyl-HiC data and bulk Methyl-HiC. Contact matrixes were normalized by sequencing coverage. c) Comparison of average global methylation level from single cell Methyl-HiC between primed and naïve mESCs. d) DNA methylation concordance on loop anchors is validated in single cell Methyl-HiC datasets. Chromatin loops detected are from bulk Methyl-HiC by HiCCUPS. e) Late replication regions defined by Repli-ChIP show significantly higher DNA methylation variation. SD means standard deviation. f) Late replication regions defined by Repli-ChIP show significantly lower variation in chromatin contact. SD means standard diversion.

The single cell Methyl-HiC dataset would enable analysis of cell-to-cell variability in both chromosome organization and DNA methylation as they relate to different replication timing regions. DNA replication is accompanied by dynamic chromosome organization and DNA methylation. In particular, previous studies have uncovered a link between chromatin compartments and timing of DNA replications, and between topologically associating domains with DNA replication domains^27,28^. While DNA methylation pattern is generally replicated by the DNMT1 at each cell division, the fidelity is not 100%, resulting in variation and plasticity in the methylome^29^. We partitioned the genome into four groups based on replication timing previously determined by Repli-ChIP data from mESC^30^. We then analyzed single cell methylomes and 3D interactions in each replication timing group. We found that regions associated with late replication showed higher DNA methylation levels (Supplementary Fig. 3f), consistent with their heterochromatin nature^31^. Moreover, the late replication regions tend to show higher cell-to-cell variability of DNA methylation evidenced by increased Standard Deviation (SD) across 107 mESCs (Fig. 3e) and a higher rate of hemi-methylation (CpGs with methylation level equal to 0.5 in single cell DNA methylome) (Supplementary Fig. 3g). Interestingly, late-replicating DNA had lower variabilities in chromatin interactions than early-replicating DNA (Fig. 3f). Taken together, our data suggest that early replication regions are characterized by more dynamic long-range interactions, less variable DNA methylation, more permissive chromatin states and higher gene expression level^32^, than late replication time domains.

Epigenetic heterogeneity in tissues or cell population presents a significant challenge in analysis of epigenome of tissue samples^17,33^. As DNA methylation differs among different cell types, it is possible to deconvolute distinct cell types from single cell Methyl-HiC datasets based on DNA methylation so that the chromatin contacts could be investigated in each cell type. Indeed, the primed and naïve mESCs were clearly separable using single cell Methyl-HiC datasets (Fig. 4a). Interestingly, the primed mESCs can be further divided into two subpopulations according to DNA methylome, which is consistent with previous report^18^ (Fig. 4a). We then compared these subpopulations with lineage specific DNA methylation profiles from ENCODE projects and found that the two subpopulations in primed cells clustered with different lineages specific features, in which Cluster 3 showed potential embryonic limb development trend (Fig. 4b). We then aggregated the contact matrix of cells that share the same DNA methylation status, and compared the populations to each other as well as clearly defined mESCs cells. Our result showed that the aggregated contact matrix could show heterogeneity between different DNA methylation clustered cells (Fig. 4c), which suggests that our method will be useful to determine single cell HiC identity of heterogeneous cell population and tissues. To further show the biological function of the 3D structure heterogeneity, we identified the differential compartments between two clusters (Supplementary Table 3). Genome ontology (GO) analysis of the Differential Methylated Regions (DMRs) and genes located in these differential compartments revealed that DMRs and genes that switched from compartment B in Cluster 2 to compartment A in Cluster 3 were enriched for genes with function related to embryonic limb development (Fig. 4d and 4e). For example, *HoxD* cluster genes and *Epha4* are key regulators for embryonic limb development^34,35^. Here we showed that *HoxD* cluster genes and *Epha4* gene switched from compartment B in Cluster 2 to compartment A in Cluster 3 (Fig. 4f and Supplementary Fig. 4). This is in line with our above clustering according to DNA methylome, indicating that our method can discover differential chromatin conformations in heterogeneous cells.

**Fig.4.**
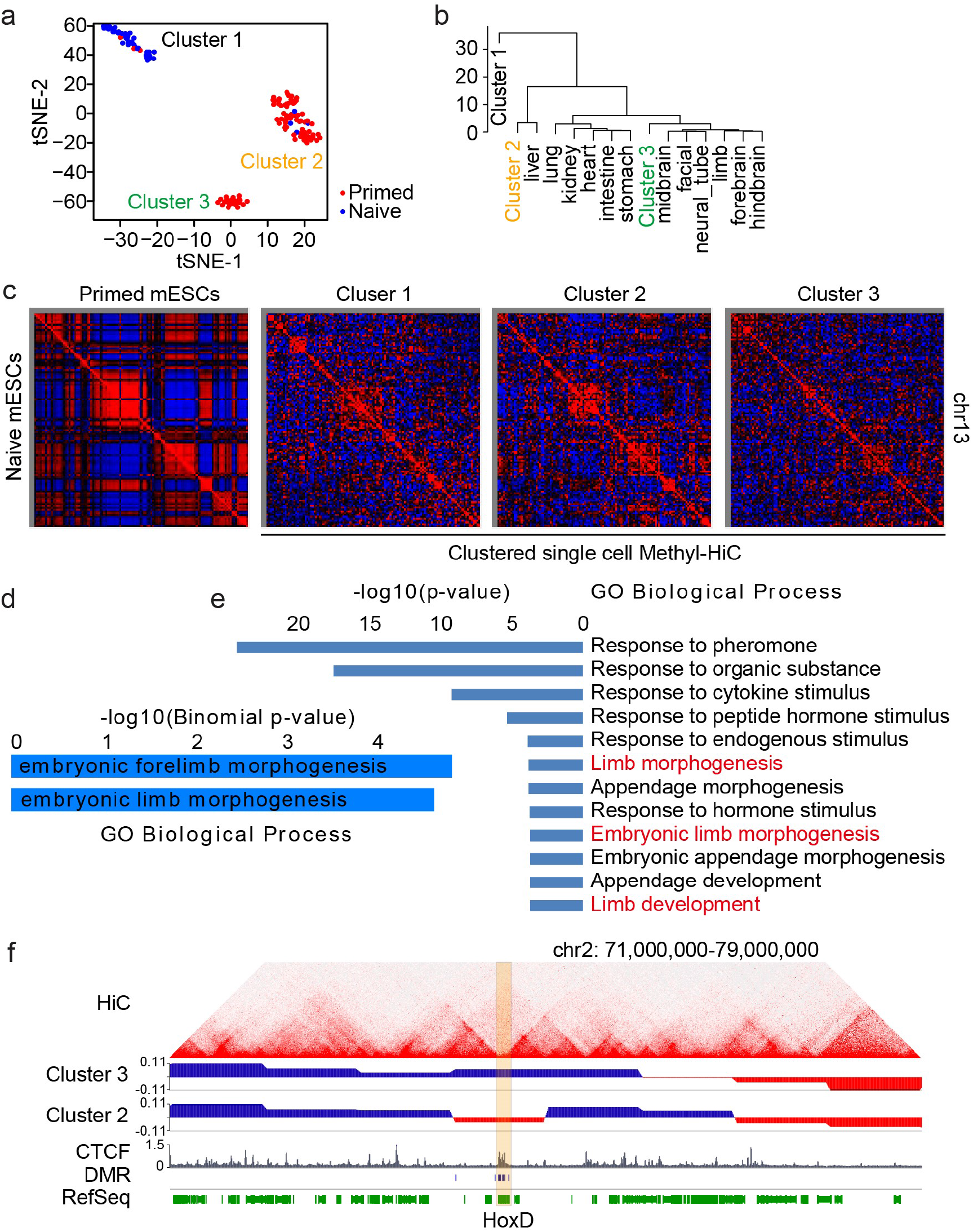
Single cell Methyl-HiC reveals heterogeneity of cultured mouse embryonic stem cells. a) T-sne visualization of unsupervised clustering results according to DNA methylation. Methylation level is calculated in non-overlaping 1Mb bins. b) Subgroups in primed cells show lineage specific DNA methylation. Cells from each cluster are aggregated and compared with tissue-specific methylome. c) Pearson correlation matrixes of contact matrix from different cell clusters. The left map shows the bulk *in situ* Hi-C matrixes from primed and naïve cells, respectively. Pearson correlation is calculated under 1Mb resolution. Color ranges have been set to the same scale. d) GO biological process terms for DMRs in Compartment B in Cluster 2 while switch to Compartment A in Cluster 3. e) GO biological process terms of genes switching from Compartment B in Cluster 2 to Compartment A in Cluster 3 f) Snapshot of *HoxD* cluster genes in different compartments between Cluster 2 and 3.

Here, we report a novel method, Methyl-HiC, which combines high throughput chromatin conformation capture with whole genome bisulfite sequencing that can simultaneously profile and integrate two epigenetic regulations on the same DNA molecule. We demonstrate that Methyl-HiC could be used to study the higher-order organization of chromosomal structures and DNA methylomes simultaneously in mixed populations and in single cells. By doing these, we show that DNA methylation status is generally concordant between spatially close cytosines.

The simultaneous profiling of multiple epigenomic features, especially from single cells, has multiple advantages over assaying each feature individually. As the cost of both Hi-C and WGBS is predominantly sequencing, combining two assays together will lead to substantial savings due to reduced sequencing cost. In addition, Methyl-HiC is desirable when the biological material is limited, such as oocytes, zygotes and early embryos. In particular, single cell Methyl-HiC generates maps of DNA methylome and chromosome conformation from the same cells, allowing integrative analysis of chromatin architecture and epigenome in individual cells. While current protocol of single cell Methyl-HiC is still limited by data sparsity, this limitation could be overcome in the future by better DNA methylation detection strategies, such as bisulfite free DNA methylation profiling^36^, or better amplification approaches, such as META (multiplex end-tagging amplification)^37^. Further development of single cell Methyl-HiC in the future to include variants detection from bisulfite reads^38^ and transcriptome^39,40^ from the same cell will allow us to reveal the comprehensive regulation diagram in a single cell from genetic, epigenetic, transcription, and chromatin interaction aspects.

Although single cell epigenomic datasets can be clustered *de novo*, the identities of each sub-population are difficult to determine because of the stochasticity of epigenomic regulation and lack of reference epigenome from previously characterized cell types, especially for assays like single cell Hi-C that the reference of rare populations are very hard to profile. DNA methylation is a stable and cell-type specific epigenetic modification, which has proved to be capable for analyzing the cell composition of heterogeneous samples, such as neuronal subtypes in cortex^17^. Thus, by combining single cell DNA methylation with single cell Hi-C, chromosome conformation heterogeneity can be revealed by grouping via DNA methylation. Because of its low cell-input requirements, single cell Methyl-HiC is readily applicable to diverse samples, tissue types, and rare cell populations, which will benefit our understanding of the chromosome conformation in different conditions.

## Methods

### Cell culture

The F1 Mus musculus castaneus × S129/SvJae mouse ESC line (F123) was a gift from the laboratory of E. Heard and has been described previously^41^. Cells in primed state were cultured with irradiated mouse embryonic fibroblasts (Gibco, A34180) in medium with 85% DMEM, 15% Knock-out Serum Replacement (Gibco, 10828-028), 1X penicillin/streptomycin, 1X non-essential amino acids (Gibco, 11140-050), 1X GlutaMax (Gibco, 35050), 0.4 mM β-mercaptoethanol and 1000U/ml LIF (Millipore, ESG1107). Cells in naïve state were adapted from primed cells by passaging cells in MEF free and serum free conditions in 2i medium, which contained 50% NEUROBASAL (Gibco 21103-049), 50% DMEM/F12 (Gibco 11320-033), 0.5% N2-SUPPLEMENT (Gibco 17502-048), 1% B27+RA (Gibco 17504-044), 0.05% BSA (Gibco 15260-037), 1X penicillin/streptomycin, 2mM GLUTAMINE (Gibco 25030-081), 150 μ M Monothioglycerol (Sigma M6145), 1000U/ml LIF (Millipore, ESG1107), 1 μ M MEK inhibitor (Stemgent 04-0006), and 3 μ M GSK3 inhibitor (Stemgent 04-0004). Primed cells were collected and plated on 0.2% gelatin-coated feeder-free plates for 30 mins before harvesting to remove feeder cell contamination.

### Methyl-HiC

*In situ* Hi-C was performed according to previous protocol^12^. Briefly, two million cells were cross-linked with 1% formaldehyde for 10 min at room temperature. Reaction was then quenched with 0.2M Glycine. Cell pellets were washed with cold PBS and lysed with lysis buffer to get nuclei pellets. Nucleus were permeabilized with 0.5% SDS. DNA was *in situ* digested with 100 units of DpnII (NEB) overnight. The ends of restriction fragments were filled with biotinylated nucleotides and *in situ* ligated. After reversal of crosslinks, ligated DNA was ethanol precipitate and sheared to a length of ~400bp by sonication (Covaris). Sonicated products were pulled down with streptavidin beads. Library construction was then performed on beads. After adapter ligation, beads were suspended in 20ul TE buffer and subjected to bisulfite conversion with EZ DNA Methylation-Gold™ Kit (Zymo, D5005). Unmethylated lambda DNA was sonicated and ligated with the same adapter for Methyl-HiC sample and then was spiked in at 0.5% before bisulfite conversion. After conversion, streptavidin beads were removed with magnet and the supernatant were purified. Purified bisulfite converted DNA was amplified with HiFi Hotstart Uracil+Ready Mix (KAPA, KK2802).

### Single cell Methyl-HiC (scMethyl-HiC)

*In situ* Hi-C was performed as same as above bulk Methyl-HiC till proximal ligation. After ligation, nuclei pellets were centrifuged and washed with PBS. Pellets were suspended in PBS stained with 1:200 DRAQ7 (CST, 7406S). FACS sorted single nuclei were directly added to 96 well plate with 9ul PBS in each well. Sorted nuclei were then briefly centrifuged and bisulfite conversion was directly performed on the sorted nuclei with EZ-96 DNA Methylation-Direct^™^ Kit according to the manufactory manual (Zymo, D5020). 0.5% of fragmented lambda DNA was spiked in before bisulfite conversion. Following bisulfite conversion of single nuclei, random priming of bisulfite-converted DNA with high concentration Klenow fragments (Enzymatics, P706L) incorporates an indexed P5 adapter to 5’ ends of synthesized fragments, which can be used for downstream multiplexing capability. Exonuclease I and Shrimp Alkaline Phosphatase (NEB) treatments were then performed to digest unused random primer and inactivate dNTPs, followed by a SPRI bead-based purification step. P7 adapter were then ligated to the 3’ end of single-stranded products by Adaptase module (Swift, 33096)^42^. Library amplification was then performed using indexed primers that incorporate dual indexing to enable 96-plex sequencing. Amplified libraries were pooled together and subjected to size selection and library quantification.

### Whole-genome bisulfite sequencing

Genomic DNA was first extracted from mESCs (DNeasy Blood & Tissue Kit, Qiagen). 1–1.5 μg of genomic DNA was fragmented by sonication (Covaris), end-repaired, dA-tailed and ligated to cytosine methylated Illumina Truseq adapter. Ligation product was subjected to bisulfite conversion reaction according to the manufacturer’s instructions (EZ DNA Methylation-Gold Kit, Zymo, D5005). 0.5% unmethylated λDNA was spiked-in before the conversion. Bisulfite converted DNA was then PCR amplified and purified.

### Sequencing of DNA libraries

The quantification of sequencing libraries was determined by qPCR and TapeStation DNA analyzer (Agilent Technologies). Pooling of multiplexed sequencing samples, clustering and sequencing were carried out as recommended by the manufacturer on Illumina HiSeq 2500 or HiSeq 4000. For bisulfite-converted libraries, at least 50% of balanced libraries or Phix were multiplexed to overcome the imbalance of GC ratio. All libraries were sequenced in paired-end mode.

### Analysis of methylation data

Raw reads were first trimmed as paired-end reads using Trimmomatic with default parameters to remove the adapters and low quality reads. Trimmed reads were aligned to mm9 using Bismark (v12.5). PCR duplications were removed with Picard (http://broadinstitute.github.io/picard/). CpG methylation level were calculated by Bis-SNP in bissnp_easy_usage.pl with default parameters.

### Analysis of Hi-C data

All sequence data were produced using Illumina paired-end sequencing. Each end of the raw reads was mapped separately to the mm9 reference genome using BWA-mem. Filtered reads were then paired and de-duplicated (Picard). Reads that map to the same fragment were further removed. Contact matrices were generated at different resolution using Juicer pipeline with KR normalization and visualized using Juicebox. Loops were then called by HICCUPS.

### Methyl-HiC reads mapping by Bhmem

Raw reads were first trimmed as paired-end reads using Trimmomatic with default parameters to remove the adapters and low-quality reads. All C in mm9 reference genome are converted to T to make C_to_T reference genome, and all G are converted to A to make G_to_A reference genome. On paired-end reads, C is converted to T in end 1 and G is converted to A in end 2. The converted end 1 and end 2 are joined as single paired fragment when both of them were mapped to C_to_T reference genome or G_to_A reference genome by BWA-MEM. Only unique mapped and passed VendorQualityFilter reads on both ends were joined. The best paired reads were selected based on the following priorities: (1) mapping quality on both ends are bigger than 0. (2) the sum of mapping quality score is larger. (3) both ends of reads are mapped into the same chromosome. (4) the sum of alignment score is larger. (5) the sum of mismatches is smaller. (6) the sum of matched cigar string length is larger. Only best paired reads were output for the following analysis. Low quality reads (not unique mapped on both ends, PCR duplicate and mapping quality score < 30) were removed for the following downstream analysis. Only bases with quality score more than 5 were included in the downstream methylation analysis. Details were implemented in Bhmem.java.

### Methylation concordance analysis in Methyl-HiC

Bisulfite incompletely converted reads and 5’ end incompletely converted cytosine (M-bias) were filtered out as that in Bis-SNP^38^. Only read pairs with methylation level at both ends were kept for the methylation concordance analysis. For the regions of interests, such as HiCCUPS loops anchor regions, Pearson correlation coefficient (PCC) was calculated by the methylation level at each end of paired reads rather than by the average methylation level at each end of paired regions. Only reads with at least 20kb distant from each end were considered for the long-range methylation concordance. Expected control was generated by shuffling the read pairs within the same genomic regions. Fisher z-transformation, implemented in function of “cocor.indep.groups” at R package “cocor” was used to determine the significance between observed PCC and control PCC. Details were implemented in MethyCorAcrossHiccups.java.

### Single cell Methyl-HiC analysis

Reads were mapped by the same Bhmem pipeline with additional parameters to adapt the single cell protocol (-nonDirectional -pbat, G is converted to A in end 1 and C is converted to T in end 2; End 1 and end 2 mapped to different bisulfite converted genomes were also considered as the candidate of best pair when join them). We also utilize the restriction enzyme cutting sites on the reference genome to help the reads mapping. Only read pairs with restriction enzyme cutting sites nearby the reads (+/-50bp nearby alignment start and end of the reads) were kept for the further analysis.

After removing low quality reads (not unique mapped, PCR duplicate, not both ends unique mapped and mapping quality score < 30), cells with less than 250,000 reads and 10,000 ligation events were removed for the further analysis. Finally, there are 47 naïve cells and 103 primed cells left for all of the single cell analysis.

Methylation density (total_methylated_count/total_count) was calculated in each non-overlapped 1Mb window at mm9 autosome chromosomes. Missing data was replaced by the mean methylation level at the same bin across all single cells. Bins with missing data in all of cells were removed from analysis. Then t-Distributed Stochastic Neighbor Embedding (t-SNE) was applied to represent the methylation level structure in single cell level (Rtsne package in R 3.4.0 with plexity=10 and max_iter=5000). Different plexity level and random seed were applied to test the robustness of representation. Methylated count and total count (missing data is not replaced by mean value yet) at each window within each sub-cluster was aggregated and then calculated the methylation level. Methylation level at each CpG from external ENCODE methylomes were liftovered from mm10 to mm9. Overlapped CpGs after liftover were discarded. Methylation density was calculated in each non-overlapped 1Mb window for ENCODE data. Bins with missing data in all of samples were removed from analysis. Euclidean distance between 3 merged sub-clusters in single cell Methyl-HiC and external ENCODE methylomes was calculated and clustered by “ward.D2” by hclust function in R. BAM files were merged within each sub-cluster and then visualized for Hi-C contact frequency by Juicebox (1.9.0)

### Replication timing

Wavelet-smoothed of mean late/early S-phase ratios data by Repli-ChIP is obtained from ENCODE in ES-46C cell line. Regions with high values indicate domains of early replication where initiation occurs earlier in S-phase or early in a higher proportion of cells.

## Data availability

Figures show merged data from all replicates. The Methyl-HiC, single cell Methyl-HiC, and WGBS data sets generated in this study have been deposited in the Gene Expression Omnibus (GEO) under the accession number GSE119171. Previously published data used for this study are listed in Supplementary Table 4. All the source code for Methyl-HiC analysis is publicly available at Bitbucket: https://bitbucket.org/dnaase/bisulfitehic/src/master/

## Acknowledgments

We thank Samantha Kuan and Dr. Bin Li for processing of sequencing. We also thank Dr. Ramya Raviram for kindly revision and comments on the manuscript. Funding from the NIH 4D Nucleome program (U54DK107977) supported the project. G.L. and B.R. were supported by NIH 4D Nucleome program (U54DK107977). Y.L. and M.K. were supported by funding from NIH 1U01HG007610-01.

## Author contributions

B.R., Y.L., and G.L. conceived the study and prepared the manuscript. G.L. performed the Methyl-HiC and single cell Methyl-HiC experiments. Y.L. wrote new Methyl-HiC analysis pipeline and performed the analysis. Y.L. and G.L. performed the *in situ* Hi-C and WGBS analysis. G.L. and Y.L. wrote the manuscripts with the guidance of M.K. and B.R. Correspondence and requests for materials should be addressed to B.R.

## Supplementary Materials

**Fig. S1.**
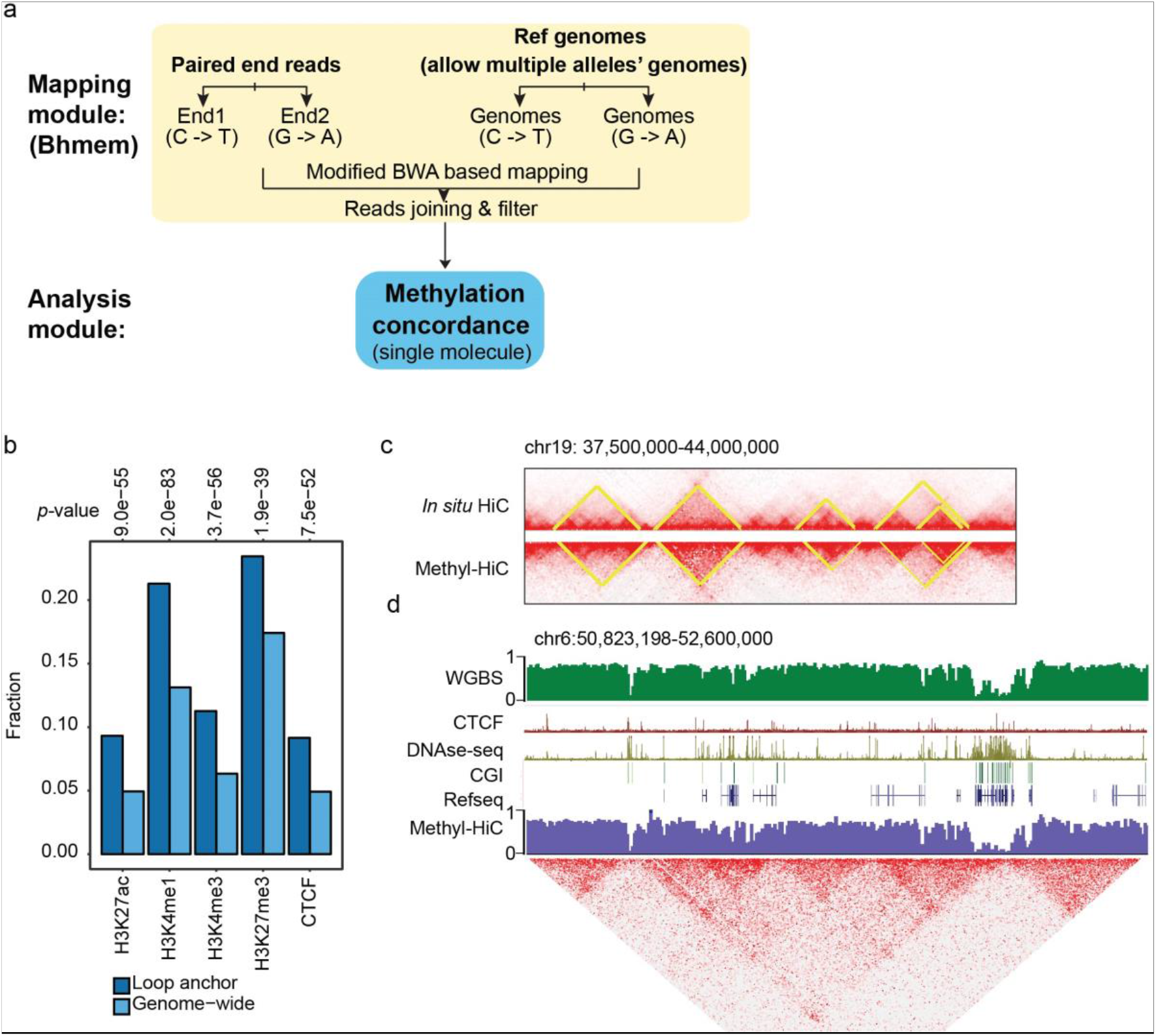
Methyl-HiC simultaneously profiles long-range chromatin interactions and DNA methylome in mouse embryonic stem cells. a. The Bhmem Methyl-HiC data mapping and analysis modules. b. HiCCUPS loops called from Methyl-HiC dataset enrich enhancers and promoters with active histone markers, Polycomb-Repressed chromatin marked by H3k27me3, and CTCF. c. Contact domains called from *in situ* HiC and Methyl-HiC, respectively d. A snapshot of both DNA methylation and HiC contact matrix

**Fig. S2.**
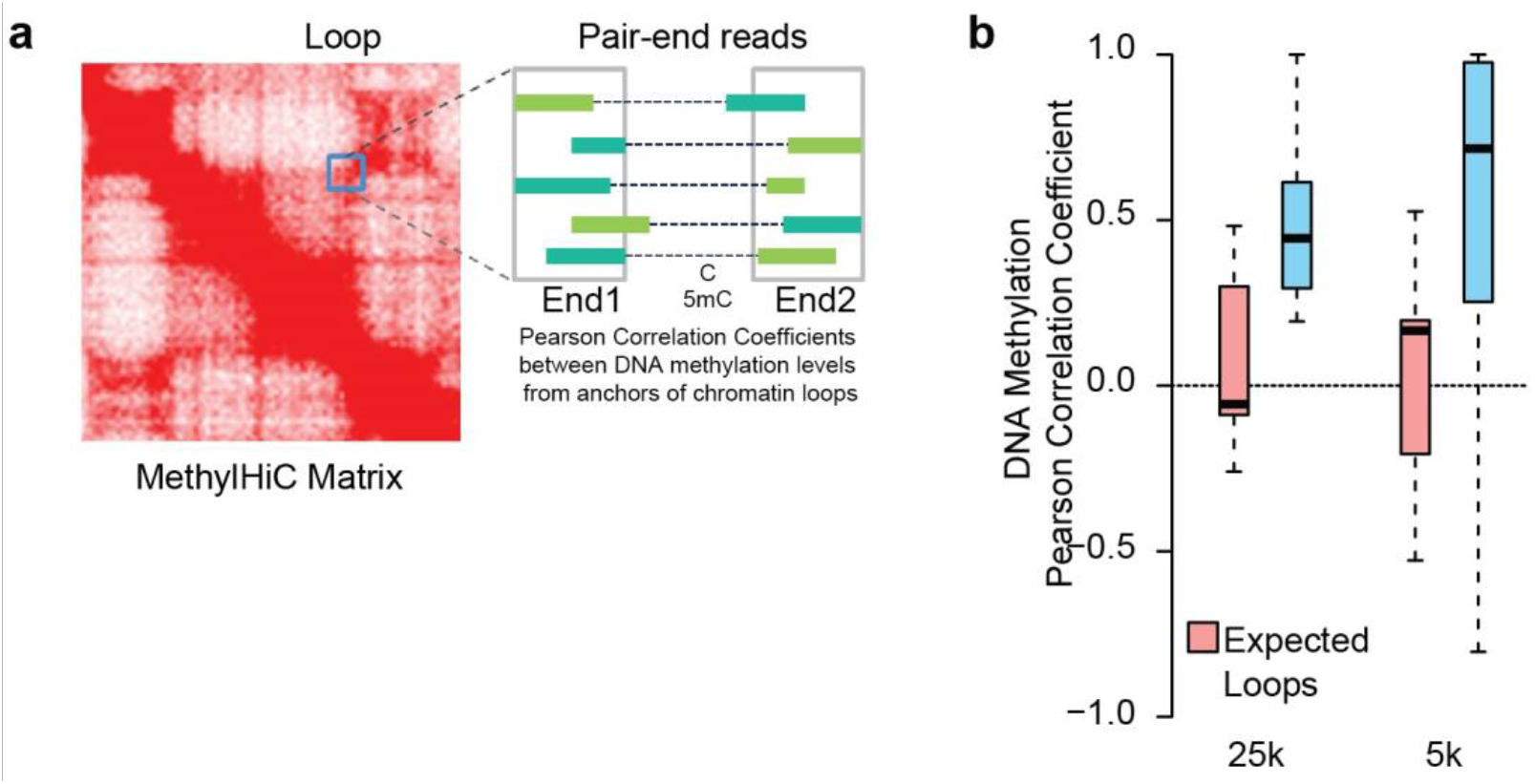
DNA methylation status is generally concordant between spatially proximal regions. a. Illustration of the calculation for methylation concordance on DNA loop anchors b. Pearson correlation coefficients distribution of individual loops at two different resolutions.

**Fig. S3.**
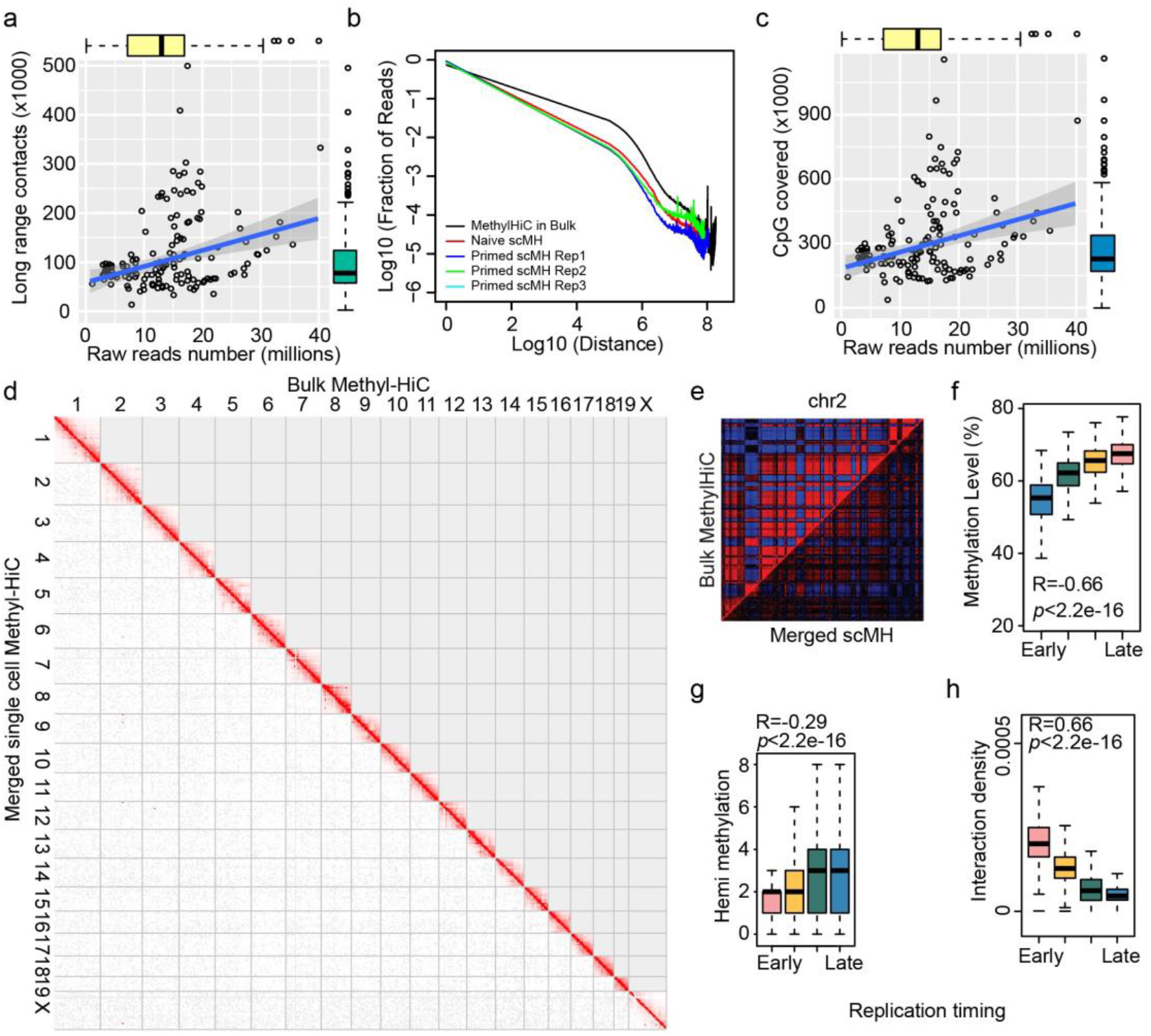
Simultaneous analysis of DNA methylome and chromatin architecture in individual cells by Single Cell Methyl-HiC. a. The scatterplot for raw reads numbers and the number of CpG covered. b. The scatterplot for raw reads numbers and the number of long range contacts. c. Contact probability of merged single cell Methyl-HiC and bulk Methyl-HiC data. d. Genome wide contact matrix comparison between resembled single cell Methyl-HiC data and bulk Methyl-HiC data. e. Pearson correlation matrix comparison between ensemble single cell Methyl-HiC data and bulk Methyl-HiC data. f. DNA methylation distribution in replication timing regions. g. The hemimethylation density in replication timing regions. h. Interaction density distribution in replication timing regions.

**Fig. S4.**
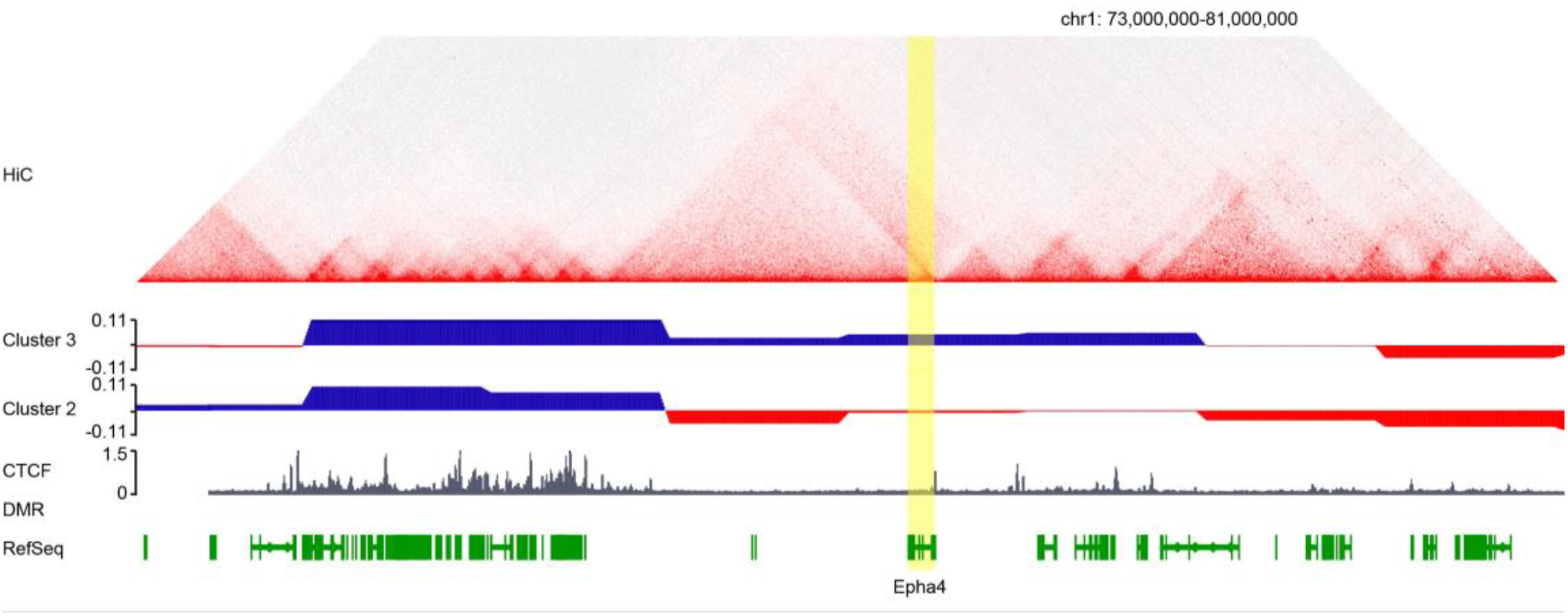
Single cell Methyl-HiC reveals heterogeneity of cultured mouse embryonic stem cells. The snapshot for *Epha4* gene in different compartments between Cluster 2 and Cluster 3.

**Supplemental Table S2.**
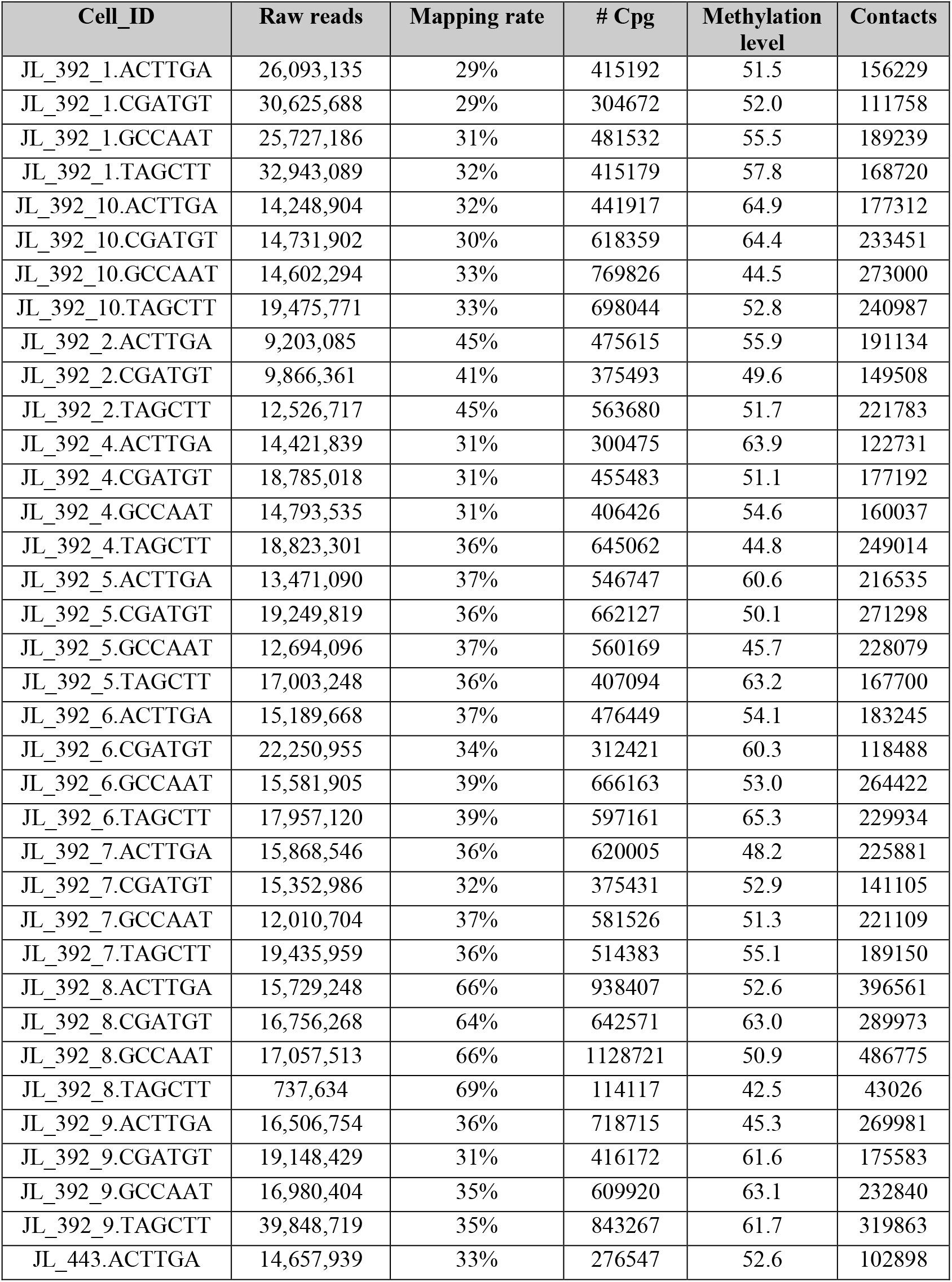
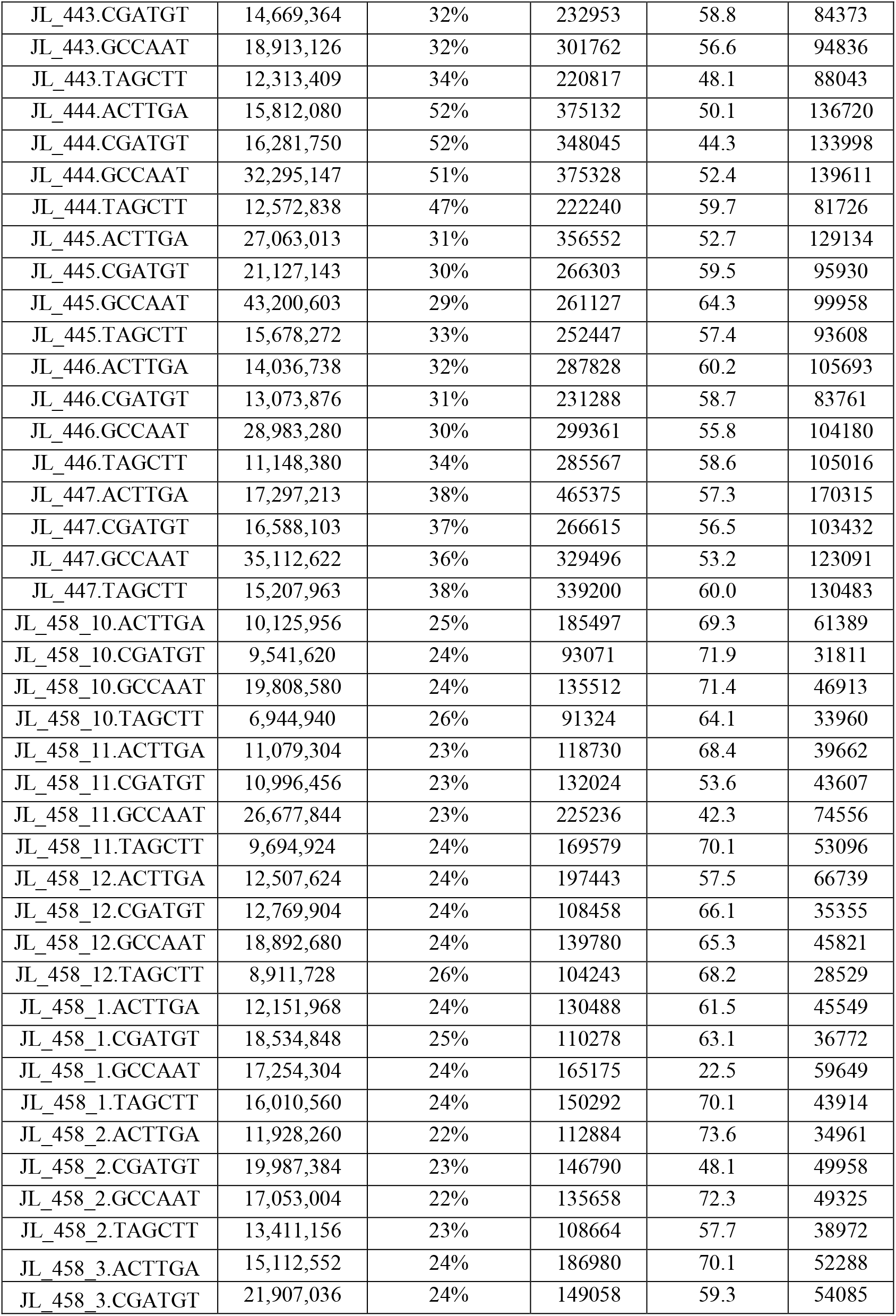
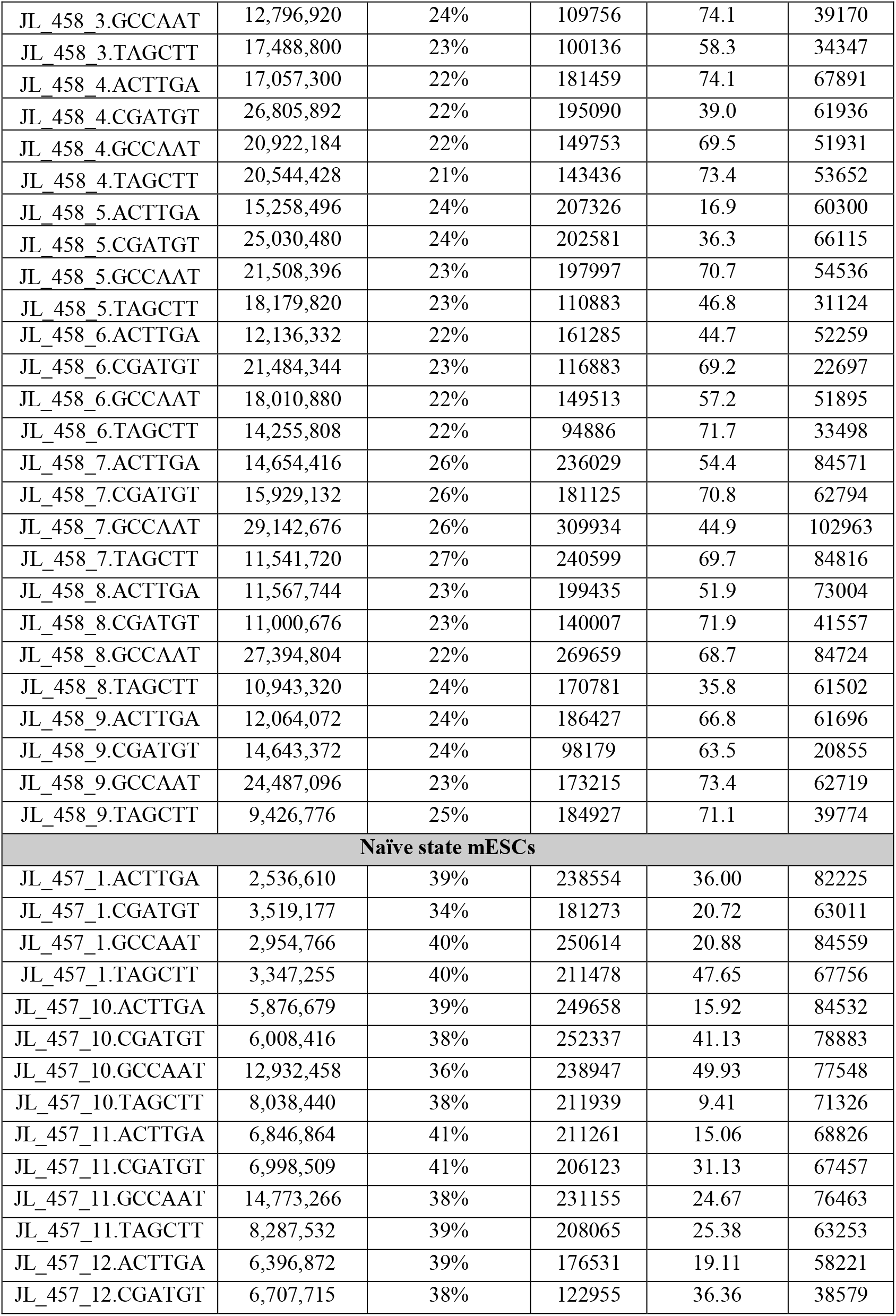
Summary of single cell Methyl-HiC datasets.

**Supplemental Table S3.**
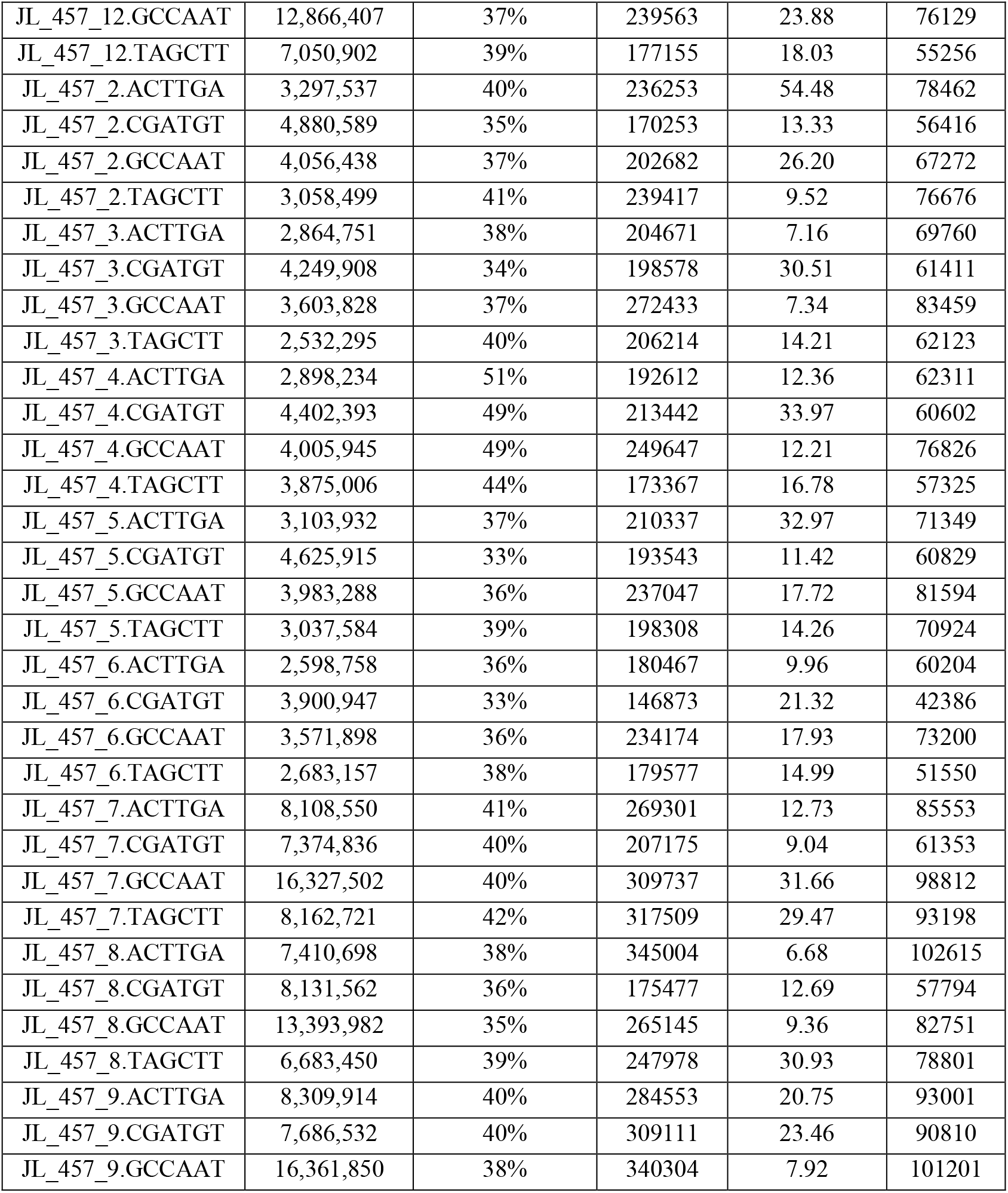
Differential Compartments between Cluster 2 and 3 in single cell Methyl-HiC.

**Supplemental Table S4.**

| Published data sets used in this study | Data Source | Reference |
| --- | --- | --- |
| F123 *in situ* HiC | GSE86150 | <sup>1</sup> |
| sNMseq | GSE97179 | <sup>2</sup> |
| Chromatin states in mESCs | ENCODE | <sup>3</sup> |
| Single cell Methylome in primary and naïve mESCs | GSE56879 | <sup>4</sup> |

## References

1. Iurlaro, M., von Meyenn, F. & Reik, W. DNA methylation homeostasis in human and mouse development. Current Opinion in Genetics & Development 43, 101–109 (2017).

2. Guo, J.U., Su, Y., Zhong, C., Ming, G.-l. & Song, H. Hydroxylation of 5-methylcytosine by TET1 promotes active DNA demethylation in the adult brain. Cell 145, 423–434 (2011).

3. Lyko, F. The DNA methyltransferase family: a versatile toolkit for epigenetic regulation. Nat Rev Genet 19, 81–92 (2018).

4. Guo, S. et al. Identification of methylation haplotype blocks aids in deconvolution of heterogeneous tissue samples and tumor tissue-of-origin mapping from plasma DNA. Nature Genetics 49, 635 (2017).

5. Shoemaker, R., Deng, J., Wang, W. & Zhang, K. Allele-specific methylation is prevalent and is contributed by CpG-SNPs in the human genome. Genome Research 20, 883–889 (2010).

6. Lister, R. et al. Human DNA methylomes at base resolution show widespread epigenomic differences. Nature 462, 315 (2009).

7. Flusberg, B.A. et al. Direct detection of DNA methylation during single-molecule, real-time sequencing. Nature Methods 7, 461 (2010).

8. Rand, A.C. et al. Mapping DNA methylation with high-throughput nanopore sequencing. Nature Methods 14, 411 (2017).

9. Dixon, Jesse R., Gorkin, David U. & Ren, B. Chromatin Domains: The Unit of Chromosome Organization. Molecular Cell 62, 668–680 (2016).

10. Schmitt, A.D., Hu, M. & Ren, B. Genome-wide mapping and analysis of chromosome architecture. Nature Reviews Molecular Cell Biology 17, 743 (2016).

11. Lieberman-Aiden, E. et al. Comprehensive Mapping of Long-Range Interactions Reveals Folding Principles of the Human Genome. Science 326, 289 (2009).

12. Rao, Suhas S.P. et al. A 3D Map of the Human Genome at Kilobase Resolution Reveals Principles of Chromatin Looping. Cell 159, 1665–1680 (2014).

13. Yang, T. et al. HiCRep: assessing the reproducibility of Hi-C data using a stratum- adjusted correlation coefficient. Genome Research (2017).

14. Fortin, J.-P. & Hansen, K.D. Reconstructing A/B compartments as revealed by Hi-C using long-range correlations in epigenetic data. Genome Biology 16, 180 (2015).

15. Yue, F. et al. A comparative encyclopedia of DNA elements in the mouse genome. Nature 515, 355 (2014).

16. Jones, P.A. Functions of DNA methylation: islands, start sites, gene bodies and beyond. Nature Reviews Genetics 13, 484 (2012).

17. Luo, C. et al. Single-cell methylomes identify neuronal subtypes and regulatory elements in mammalian cortex. Science 357, 600 (2017).

18. Smallwood, S.A. et al. Single-cell genome-wide bisulfite sequencing for assessing epigenetic heterogeneity. Nature Methods 11, 817 (2014).

19. Guo, H. et al. Single-cell methylome landscapes of mouse embryonic stem cells and early embryos analyzed using reduced representation bisulfite sequencing. Genome Research 23, 2126–2135 (2013).

20. Nagano, T. et al. Single-cell Hi-C reveals cell-to-cell variability in chromosome structure. Nature 502, 59–64 (2013).

21. Stevens, T.J. et al. 3D structures of individual mammalian genomes studied by single-cell Hi-C. Nature 544, 59 (2017).

22. Ramani, V. et al. Massively multiplex single-cell Hi-C. Nat Methods 14, 263–266 (2017).

23. Nagano, T. et al. Cell-cycle dynamics of chromosomal organization at single-cell resolution. Nature 547, 61–67 (2017).

24. Flyamer, I.M. et al. Single-nucleus Hi-C reveals unique chromatin reorganization at oocyte-to-zygote transition. Nature 544, 110 (2017).

25. Brons, I.G.M. et al. Derivation of pluripotent epiblast stem cells from mammalian embryos. Nature 448, 191 (2007).

26. Durand, N.C. et al. Juicebox Provides a Visualization System for Hi-C Contact Maps with Unlimited Zoom. Cell systems 3, 99–101 (2016).

27. Rhind, N. & Gilbert, D.M. DNA Replication Timing. Cold Spring Harbor perspectives in biology 5, a010132–a010132 (2013).

28. Pope, B.D. et al. Topologically associating domains are stable units of replication-timing regulation. Nature 515, 402 (2014).

29. Suzuki, M. et al. Late-replicating heterochromatin is characterized by decreased cytosine methylation in the human genome. Genome Research 21, 1833–1840 (2011).

30. Hiratani, I. et al. Global reorganization of replication domains during embryonic stem cell differentiation. PLoS Biol 6, 0060245 (2008).

31. Gilbert, D.M. Replication timing and transcriptional control: beyond cause and effect. Current Opinion in Cell Biology 14, 377–383 (2002).

32. Rivera-Mulia, J.C. et al. Allele-specific control of replication timing and genome organization during development. Genome Research 28, 800–811 (2018).

33. Schwartzman, O. & Tanay, A. Single-cell epigenomics: techniques and emerging applications. Nature Reviews Genetics 16, 716 (2015).

34. Rodríguez-Carballo, E. et al. The HoxD cluster is a dynamic and resilient TAD boundary controlling the segregation of antagonistic regulatory landscapes. Genes & Development 31, 2264–2281 (2017).

35. Lupiáñez, Darío G. et al. Disruptions of Topological Chromatin Domains Cause Pathogenic Rewiring of Gene-Enhancer Interactions. Cell 161, 1012–1025 (2015).

36. Liu, Y. et al. Bisulfite-free, Base-resolution, and Quantitative Sequencing of Cytosine Modifications. bioRxiv (2018).

37. Tan, L., Xing, D., Chang, C.-H., Li, H. & Xie, X.S. Three-dimensional genome structures of single diploid human cells. Science 361, 924 (2018).

38. Liu, Y., Siegmund, K.D., Laird, P.W. & Berman, B.P. Bis-SNP: Combined DNA methylation and SNP calling for Bisulfite-seq data. Genome Biology 13, R61 (2012).

39. Clark, S.J. et al. scNMT-seq enables joint profiling of chromatin accessibility DNA methylation and transcription in single cells. Nature Communications 9, 781 (2018).

40. Cao, J. et al. Joint profiling of chromatin accessibility and gene expression in thousands of single cells. Science (2018).

41. Gribnau, J., Hochedlinger, K., Hata, K., Li, E. & Jaenisch, R. Asynchronous replication timing of imprinted loci is independent of DNA methylation, but consistent with differential subnuclear localization. Genes Dev 17, 759–73 (2003).

42. Stadler, M.B. et al. DNA-binding factors shape the mouse methylome at distal regulatory regions. Nature 480, 490 (2011).

## REFERENCES

1. Fang, R. et al. Mapping of long-range chromatin interactions by proximity ligation-assisted ChIP-seq. Cell Research 26, 1345 (2016).

2. Luo, C. et al. Single-cell methylomes identify neuronal subtypes and regulatory elements in mammalian cortex. Science 357, 600 (2017).

3. Yue, F. et al. A comparative encyclopedia of DNA elements in the mouse genome. Nature 515, 355 (2014).

4. Smallwood, S.A. et al. Single-cell genome-wide bisulfite sequencing for assessing epigenetic heterogeneity. Nature Methods 11, 817 (2014).

